# *Aedes aegypti* Malpighian tubules are immunologically activated following systemic Toll activation

**DOI:** 10.1101/2022.08.21.504691

**Authors:** Sarah D. Sneed, Sutopa B. Dwivedi, Cameron DiGate, Shane Denecke, Michael Povelones

**Affiliations:** Department of Pathobiology, School of Veterinary Medicine, University of Pennsylvania, Philadelphia, PA 19104, USA

**Keywords:** *Aedes aegypti*, Malpighian tubules, hemolymph, innate immunity, proteomics

## Abstract

**Background:** Canine heartworm is a widespread and potentially fatal mosquito-borne disease caused by infections with the parasitic nematode, *Dirofilaria immitis*. We have previously shown that systemic activation of the Toll immune pathway via silencing of the negative regulator *Cactus* in *Aedes aegypti* blocks parasite development in the Malpighian tubules, the mosquito renal organ. However, it was not established whether the Malpighian tubules were directly responding to Toll activation or were alternatively responding to upregulated proteins or other changes to the hemolymph driven by other tissues. Distinguishing these possibilities is crucial for developing more precise strategies to block *D. immitis* while potentially avoiding the fitness cost to the mosquito associated with *Cactus* silencing.

**Methods:** This study defines the transcriptional response Malpighian tubules and changes to the hemolymph proteome of *Ae. aegypti* after systemic Toll activation via intra-thoracic injection of ds*Cactus*.

**Results:** We found Malpighian tubules significantly increase expression of Toll pathway target genes that significantly overlapped expression changes occurring in whole mosquitoes. Additionally, we identified a significant overlap between the transcriptional response of the Malpighian tubules and proteins upregulated in the hemolymph.

**Conclusions:** Our data show that Malpighian tubules are capable of RNAi-mediated gene silencing and directly respond to ds*Cactus* treatment by upregulating canonical Toll pathway targets. Though not definitive, the strong correspondence between the Malpighian tubule transcriptional and the hemolymph proteomic responses provides evidence that the tubules may contribute to mosquito humoral immunity.

## Background

Vector-borne pathogens constitute a significant portion of the global infectious disease burden for both humans and animals [1]. Filarial nematodes transmitted by *Aedes*, *Culex*, and *Anopheles* mosquitoes are responsible for a significant proportion of this burden. There are currently 863 million people at risk for lymphatic filariasis, which is caused by infection with *Wuchereria bancrofti*, *Brugia malayi*, or *Brugia timori* [2, 3]. There are also millions of dogs, cats, and other small mammals at risk for infection with *Dirofilaria immitis*, the causative agent of heartworm, for which canines are the definitive host [4, 5, 6]. The mosquito innate immune system is an important determinant of vector-competency. Therefore, understanding how the mosquito responds to filarial infection and which tissues are possible sites of immune activation and pathogen restriction is essential. Identification of these mechanisms will ultimately lead to potential targets for novel transmission-blocking strategies. Modifying the immune system to block an invading pathogen without imposing a direct fitness cost on the mosquito is the foundation of many translational strategies seeking to reduce vector-borne disease.

Principal cells of the Malpighian tubules (MT), the mosquito renal organ, are invaded by *D. immitis* microfilariae (mf) which are acquired in the blood meal of a female mosquito. In the principal cells of susceptible mosquitoes (such as *Ae. aegypti* “Blackeye” strain; *Ae. aegypti*^*BE*^), microfilariae develop into L1 larvae and then molt twice forming L3 larvae which migrate through the body cavity, reside in the head and labial sheath of the proboscis, and are deposited in a drop of hemolymph on the skin of a host during blood feeding. In the refractory strain *Ae. aegypti* “Liverpool”, microfilariae invade principal cells but fail to develop to L1 [7, 8, 9, 10, 11]. This demonstrates that the MT are a critical tissue that can restrict parasite development. Comparing the transcriptional response of the MT during *D. immitis* infection, we found that both susceptible and refractory strains upregulate immune genes, but the magnitude of the response was significantly greater in the refractory strain [7]. Provoking strong immune activation by RNA interference (RNAi)-mediated gene silencing of the Toll pathway negative regulator, *Cactus*, greatly reduced the number of *D. immitis* and *B. malayi* L3 capable of emerging from susceptible mosquitoes [7, 12]. In the case of *D. immitis*, the reduction in emerging L3 could be accounted for by an increase in the number of immature larvae arrested in the MT [7].

Despite our previous work showing that *Cactus* RNAi-mediated Toll activation was sufficient to prevent the development of *D. immitis*, it was unclear whether this was the result of a MT-specific response or signals from other tissues. Here, we begin to address this question by performing mRNA sequencing of dissected MT from susceptible *Ae. aegypti*^*BE*^ strain treated with ds*Cactus* RNA and compared the response to that of whole mosquitoes, which have previously been shown to upregulate Toll pathway targets. We also performed proteomic analysis of the hemolymph to determine what proteins are upregulated following ds*Cactus* treatment. Our results suggest that in both the MT and the hemolymph, there is a robust upregulation of Toll-pathway effectors including antimicrobial peptides (AMPs), C-type lectins (CTLs), and CLIP-serine proteases (CLIPs). We hypothesize that the MT may contribute to the systemic immune response in the mosquito by secreting factors into the hemolymph.

## Methods

### Mosquito rearing and maintenance

*Aedes aegypti* Black Eye strain (*Ae. aegypti*^*BE*^, Eggs, NR-48921), provided by the NIH/National Institute of Allergy and Infectious Diseases (NIAID) Filariasis Research Reagent Resource Center for distribution by BEI Resources, NIAID, NIH. Mosquitoes were reared at 28 °C with a relative humidity of 75% on a 12-hour photoperiod. Larvae were fed a 1% suspension of liver powder (MP biomedicals) in water and fish food (Tetra variety pellets), while adults were fed 10% sucrose. Mosquitoes were housed in 20 cm^3^ cages (Bugdorm) at density of ≤ 1000 adults per cage. Using an artificial membrane feeder, mosquitoes were fed heparinized sheep blood (HemoStat Laboratories) warmed to 37 °C. All experiments were performed with adult female mosquitoes 3-7 days after eclosion.

### Gene knockdown

*Ae. aegypti Cactus* dsRNA was silenced by RNAi using standard protocols and as previously published [7]. Double-stranded RNA made from a region of *Escherichia coli* β-galactosidase (*LacZ)*, was used as a control. Fragments of *Cactus* (329 bp) and *LacZ* (541 bp) were amplified by PCR using 5’ and 3’ primers with overhangs containing T7 binding sites using iProof High-Fidelity Taq (Bio-Rad). PCR reaction products were run on an agarose gel to check for the correct amplicon, purified using the GeneJet PCR purification kit (ThermoFisher), and 1 μg was used to generate dsRNA using a HiScribe T7 RNA synthesis kit (NEB). Reaction products were purified with the GeneJET RNA Purification Kit (ThermoFisher), eluted using nuclease-free water, concentrated, with a SpeedVac to a final concentration of 3 μg/μL. Primers used for PCR in 5’ to 3’ orientation: Cactus For CGAGTCAACAGAACCCGAGCAG; Cactus Rev TGGCCCGTCAGCACCGAAAG; LacZ For AGAATCCGACGGGTTGTTACT; LacZ Rev CACCACGCTCATCGATAATTT. All primers listed have a T7 binding site (TAATACGACTCACTATAGGG) to the 5’ end. Three biological replicates were performed on separate mosquito generations. For each experiment, groups of 150 mosquitoes were anaesthetized using CO2 and injected intrathoracically with 69 nL of dsRNA (207 ng) using a Nanoject III injector (Drummond). Mosquitoes were recovered for 5 days under standard culture conditions and then processed. The number of dead mosquitoes was recorded and ~50 mosquitoes were used to prepare hemolymph samples for proteomics, and groups of 50 and 10 were used to prepare RNA samples for transcriptomics from dissected MT and intact whole mosquitoes, respectively.

### RNA-sequencing and processing

Groups of 10 whole mosquitoes and 50 MT were dissected in PBS 10 at a time and were then transferred in Trizol (Invitrogen) in a 1.5 mL rubber gasketed screw-cap tube and frozen −80 °C. Once all samples were collected from 3 independent mosquito generations, total RNA was isolated following the protocol from the manufacturer and resuspended in ultrapure water and frozen at −80 °C. RNA samples were sent to Novogene (Davis, United States) for RNA-sequencing. Briefly, mRNA was isolated using oligo d(T) magnetic beads, and cDNA libraries were generated using random hexamers. Unstranded libraries of 150 bp paired-end reads were then sequenced using the Novaseq6000 Illumina platform which generated FastQ files that were used for downstream analysis. FastQ files were first filtered for quality using the *fastP* (version # 0.20.1) program with the “–detect_adapter_for_pe” argument and implementing a minimum read length of 20 bp [13]. Trimmed reads were then mapped to the *Ae. aegypti* LVP_AGWG reference genome version 53 obtained from VectorBase/VEuPathDB [14, 15], using the HiSAT2 read mapper (version 2.2.0) [16]. Aligned reads were quantified using the featureCounts program [17], with the *Ae. aegypti* Gene Transfer Format file corresponding to genome version 53 (VectorBase). Differential expression analysis was then performed using the edgeR package (version 3.38.2) [18], using a false discovery rate (FDR) threshold of q<0.05 and a fold change threshold of log_2_ fold change (log_2_FC) >2. Raw counts were subsequently normalized by conversion into transcripts per million (TPM). Gene Set Enrichment Analysis (GSEA) was performed using Fast Gene Set Enrichment Analysis (FGSEA) package in R (version 1.22.0) [19] and using various gene sets derived from other studies as inputs (Table S1) [20, 21, 22, 23].

### Hemolymph sample preparation for Western blot and proteomic analyses

Samples of hemolymph were prepared according to our standard proboscis-clipping method [24]. Briefly, groups of approximately 50 mosquitoes were initially anesthetized with CO2 and then transferred to ice. They were aligned into rows, ventral side up and the proboscis cut with fine scissors at the midpoint. Gentle, even pressure was applied to the thorax to extract a drop of hemolymph which was collected in 5 μL of non-reducing SDS-PAGE sample buffer (Pierce) and approximately 15 mosquitoes were collected at a time, and then the sample was transferred to a 1.5 mL protein LoBind tube (Eppendorf). The sample was supplemented with additional 2x sample buffer to a concentration of 1 mosquito/μL and stored at −80 °C.

### LC-MS/MS Analyses and Data Processing

Hemolymph samples were processed for Liquid chromatography tandem mass spectrometry (LC-MS/MS) as a single broad band of an SDS-PAGE gel. Each sample band was excised, de-stained using acetonitrile, and then incubated with 10 mM dithiothreitol (DTT; GE Healthcare) at 30 °C for 1 hour. Subsequently, iodoacetamide solution (GE Healthcare) was added to a final concentration of 40 mM and the reaction was allowed to proceed at room temperature in the dark for 30 minutes. The solution was incubated with trypsin (Promega) in a 1:50 (w/w, enzyme/protein) ratio at 37 °C for 18 hours. The resulting peptides were desalted with a C-18 macro spin column (Harvard Apparatus) and then vacuum dried.

LC-MS/MS analysis was performed by the Proteomics and Metabolomics Facility at the Wistar Institute using a Q Exactive Plus or Q Exactive HF mass spectrometer (ThermoFisher Scientific) coupled with a Nano-ACQUITY UPLC system (Waters). The peptide samples were injected onto a UPLC Symmetry trap column (180 μm i.d. x 2 cm packed with 5 μm C18 resin; Waters). Tryptic peptides were separated by reversed-phase HPLC on a BEH C18 nanocapillary analytical column (75 μm i.d. x 25 cm, 1.7 μm particle size; Waters) using a 95 minute gradient formed by solvent A (0.1% formic acid in water) and solvent B (0.1% formic acid in acetonitrile). Eluted peptides were analyzed by the mass spectrometer set to repetitively scan m/z from 400 to 2000 in positive ion mode. The full MS scan was collected at 70,000 QE Plus or 60,000 QE HF resolution followed by data-dependent MS/MS scans at 17,500 QE Plus or 15,000 QE HF resolution on the 20 most abundant ions exceeding a minimum threshold of 10,000. Peptide match was set as preferred, exclude isotopes option and charge-state screening were enabled to reject unassigned and single charged ions.

Raw data was processed using MaxQuant (version 1.6.17.0) and the peptide search engine Andromeda. Using *Ae. aegypti* LVP version 52 (14,979 sequences) as a reference sequence, contaminants were filtered out and peptide sequence assignments were identified using the following parameters: precursor ion mass tolerance of 20 parts per million, and a fragment ion mass tolerance of 10 daltons. Peptides were searched using fully tryptic cleavage constraints and up to two internal cleavage sites were allowed for tryptic digestion. Fixed modifications consisted of carbamidomethylation of cysteine. Variable modifications considered were oxidation of methionine, protein N-terminal acetylation, and asparagine deamidation. All protein identifications reported were identified by MaxQuant with a false positive error rate of less than 1%. Results from MaxQuant were imported into Perseus software (Version 1.6.8, Proteome Software) for curation, label-free quantification analysis, and visualization. Data of all technical replicates were log_2_-transformed and normalized by subtracting the column median. FDRs were obtained using Target Decoy PSM selecting identifications with a p-value equal or to less than 0.05. For differential proteins analysis Two-tailed T-test and Benjamini Hochberg validation (p < 0.05 and log_2_FC change >1.5) were performed in Perseus. Protein set enrichment analysis of differentially expressed proteins was performed using R Bioconductor ‘fgsea” package (v1.22.0). Target proteins with enrichment scores above 70% and adjusted p-values <0.05 were considered for further analysis.

### Label-free Quantitation to determine differential protein expression

Label-free Quantitation (LFQ) is an MS1 quantitation approach based on peptide mass, intensity, and retention time. LFQ data were filtered according to the following criteria: (1) proteins were identified in at least two replicates per group; (2) at least one unique peptide per protein was identified; (3) the peptide false discovery rate (FDR) did not exceed 1%. Significance levels were statistically analyzed using a Two-tailed T-test. Proteins were considered differentially expressed between control and *dsCactus-*treated mosquitoes if there were both a p-value<0.05 and a fold-change ≥ 1.5 (ds*Cactus/*ds*LacZ*).

## Availability of data and materials

Enrichment in overlapping gene sets was performed using an online tool to compare gene enrichment (http://www.nemates.org/MA/progs/overlap_stats.html) based on Fisher’s exact test. Survival comparison of *Cactus* knockdown versus control (ds*LacZ*) was accomplished using Fisher’s test. All other comparisons were performed internally using the appropriate statistical packages (e.g. edgeR). Our data meet all the standards regarding the Minimum Information About a Proteomics Experiment (MIAPE), and data have been deposited to the ProteomeXchange Consortium (http://www.proteomexchange.org) via the PRIDE partner repository [25]. Gene identifiers described in this manuscript: LRIM1, AAEL012086; APL1/LRIM2, AAEL024406; TEP20, AAEL001794; Cactus, AAEL000709.

## Results

### Knockdown of Cactus triggers a Toll-like immune response in Malpighian tubules

To characterize the Malpighian tubule transcriptional response of Toll-pathway activation in *Ae. aegypti*^*BE*^, we injected groups of mosquitoes with ds*Cactus* and collected RNA from both whole insects and dissected MTs after 5 days. We additionally collected protein samples from the hemolymph (Figure 1A). Prior to dissection, we observed a significant decrease in mosquito survival to day 5 in the ds*Cactus* treated group compared to the ds*LacZ* treated control (Figure 1B). This was not unexpected, as Cactus silencing has been previously reported to incur a fitness cost [7, 26, 27].

**Figure 1:**
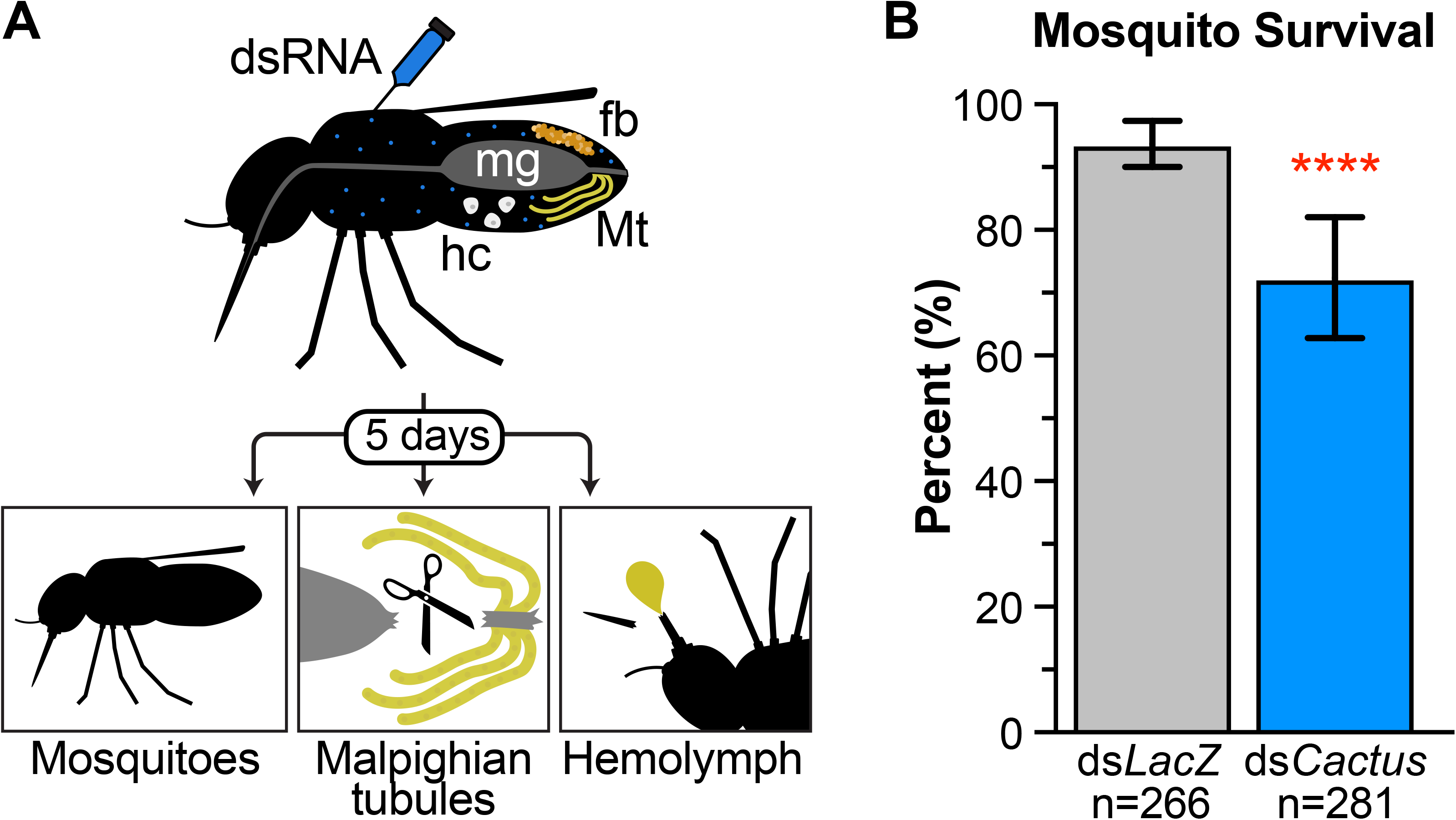
Experimental Overview. (**A**) The workflow of our experimental setup is illustrated with some mosquito tissues labeled: midgut (mg), fatbody (fb), hemocytes (hc), and Malpighian tubules (Mt). Double-stranded RNA (dsRNA) targeting the *Cactus* gene was injected into the hemocoel and five days later RNA-seq was performed on i) whole mosquitoes ii) dissected Malpighian tubules and proteomics was performed on iii) Hemolymph. (**B**) The fitness cost of ds*Cactus* treatment was estimated by scoring survival after 5 days. Significantly more individuals from the ds*Cactus* treatment (blue) died compared to the ds*LacZ* control (grey).

We performed mRNA sequencing (RNA-seq) comparing dissected MT from ds*Cactus* and ds*LacZ* treated mosquitoes as well as whole mosquitoes (Figure 1A). In total, 296,996,140 reads (average 24,749,678 per sample) were generated over a total of 12 samples (3 biological replicates each for 2 tissue types with 2 knockdown conditions) (Table S2). In all samples >90% of reads mapped to the genome and an average of 75% mapped onto annotated genes. Global variation among our samples was estimated through principal component analysis (PCA), and clustering of samples from the same biological conditions suggested strong repeatability between replicates. Differences between the MT and whole-body samples were largely based on the first principal component representing 67.8% of variation while differences between *Cactus* knockdown and control samples encompassed an additional 15.5% of variation (Figure 2A).

**Figure 2:**
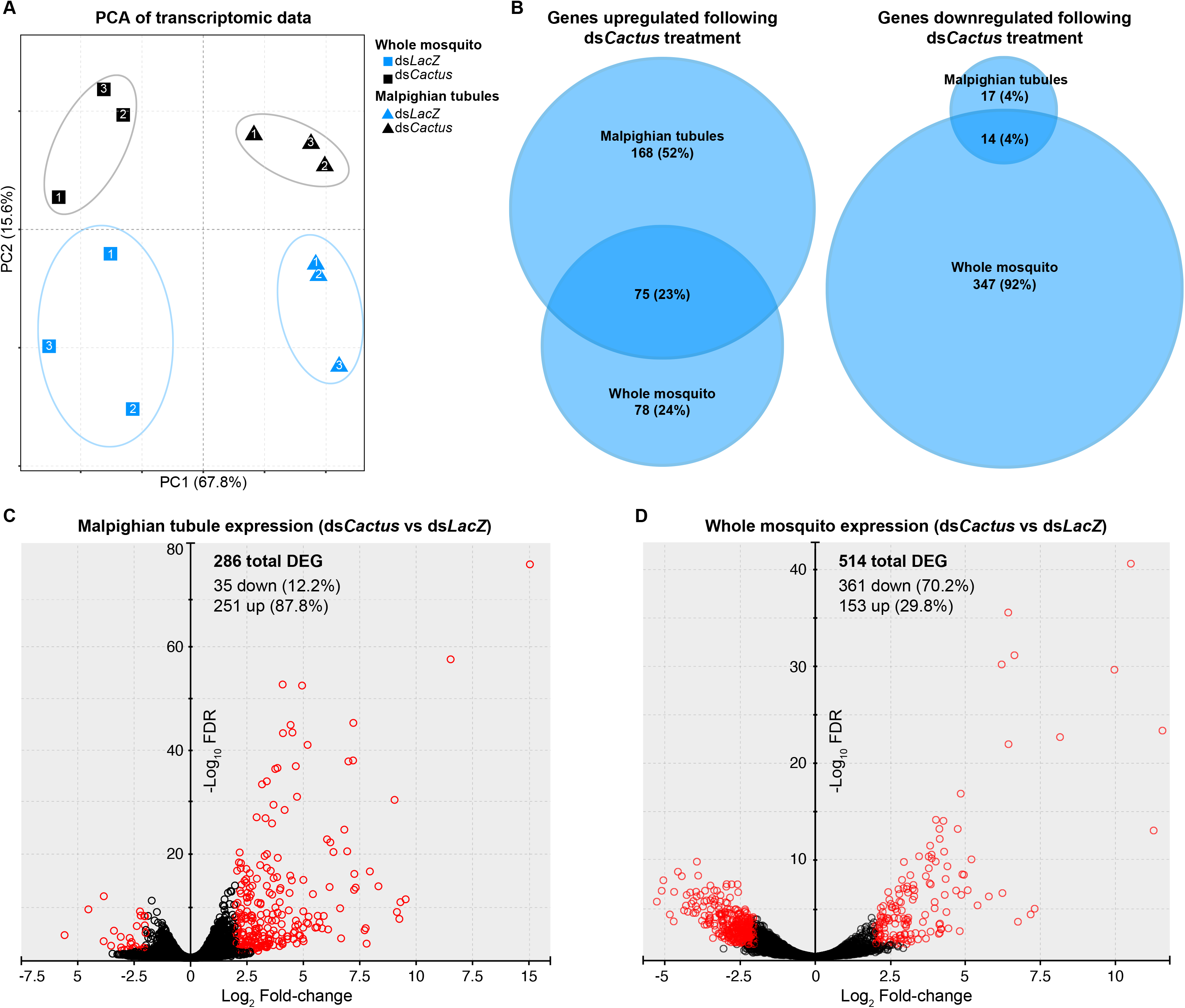
Comparisons of transcriptional responses of dissected Malpighian tubules and whole mosquitoes following Toll activation. (**A**) Principal component analysis of RNA-sequencing of whole mosquitoes (squares) and dissected Malpighian tubules (triangles) from both ds*LacZ* (blue) and ds*Cactus* (black) treatments was visualized for global variation. (**B**) Differentially expressed genes between ds*Cactus* and ds*LacZ* treatments were compared between whole bodies and Malpighian tubule specific transcriptomes. Volcano plots were made to visualize the variation in gene expression comparing ds*LacZ* versus ds*Cactus* in the (**C**) Malpighian tubules or (**D**) in whole mosquitoes. Red symbols indicated genes that significantly differentially expressed.

Differential expression (DE) analysis was performed between ds*Cactus* and ds*LacZ* transcriptomic datasets for both the MT and the whole bodies, yielding a total of 789 DE genes across the two comparisons (Table S3). A significant overlap existed between the DE gene sets of MT and whole bodies (Figure 2B; Representation factor 24.2; p < 9.045e-89). Interestingly, the vast majority of differentially expressed genes in the ds*Cactus* MT were significantly upregulated (243 genes) while fewer were downregulated (31 genes). In contrast, the whole body had a comparatively larger number of downregulated (343) vs upregulated (153) genes (Table S3; Figure 2C). This suggests that treatment with ds*Cactus* drives a physiological program of increased transcriptional activity in the MT that is proportionally “more activating,” or polarizing, than the transcriptional programs activated in the whole body in which the numbers of upregulated and downregulated genes are more similar.

Preceding formal analysis, several interesting transcripts stood out among those differentially expressed in the MT. The genes AAEL026300 and AAEL017380 showed extraordinarily high levels of abundance (>100 TPM) in *dsCactus* tubules while being almost completely absent in control tubules, showing fold changes an order of magnitude higher than any other genes in this study. Both genes encode small (~100 aa), glycine-rich secreted proteins characteristic of glycine-rich antimicrobial peptides characterized in other arthropods [28, 29, 30, 31, 32]. Among the 31 genes downregulated in *dsCactus* tubules, three were leucine rich immune protein family members (LRIMs) and two were prophenoloxidases (PPO) (Table S3). The *Cactus* gene itself was not DE in any sample, but this agrees with previous studies that have shown that genetic knockdown of *Cactus* occurs on short time scales, within the first 6-12 h post-injection, with phenotypes persisting far longer [7, 33, 34].

To elucidate relevant relationships within the genetic response to Toll activation, we performed gene set enrichment analysis (GSEA), which maps user-provided gene lists onto our Malpighian tubule transcriptomic data ordered by fold change. Certain gene families with known essential roles in immunity such as CLIP-domain serine proteases (CLIPs), C-type lectins (CTLs), anti-microbial peptides (AMPs), and serpins (SRPNs) were significantly enriched in the genes upregulated in response to ds*Cactus* treatment (Figure 3, Table S4). Interestingly, prophenoloxidases (PPOs), another immunity-associated gene family, showed the opposite association and were significantly enriched in the control (i.e. the genes downregulated in response to ds*Cactus* treatment). Subunits of the vacuolar-type ATPase (vATPase) pump were also checked given the key role of this protein in epithelial physiology [35]; these genes were also significantly downregulated in *dsCactus*-treated tubules (Table S4). We additionally considered published transcriptomic datasets reflecting genes upregulated during *B. malayi* [20], *D. immitis* [7], and *Wolbachia* [21] infection, observing that genes that were upregulated during infection were enriched in genes upregulated during ds*Cactus* treatment (Figure S1). Lastly, we observed significant correlation between genes upregulated following Toll activation via *dsCactus*-treatment and activation of this pathway driven by transgenic overexpression of REL1 and REL2, transcription factors which activate the Toll pathway [23] (Figure S1). Collectively this data suggests that in response to *dsCactus*-treatment, the MT activate canonical transcriptional signatures of immune activation that closely align with known transcriptional responses to infection.

**Figure 3:**
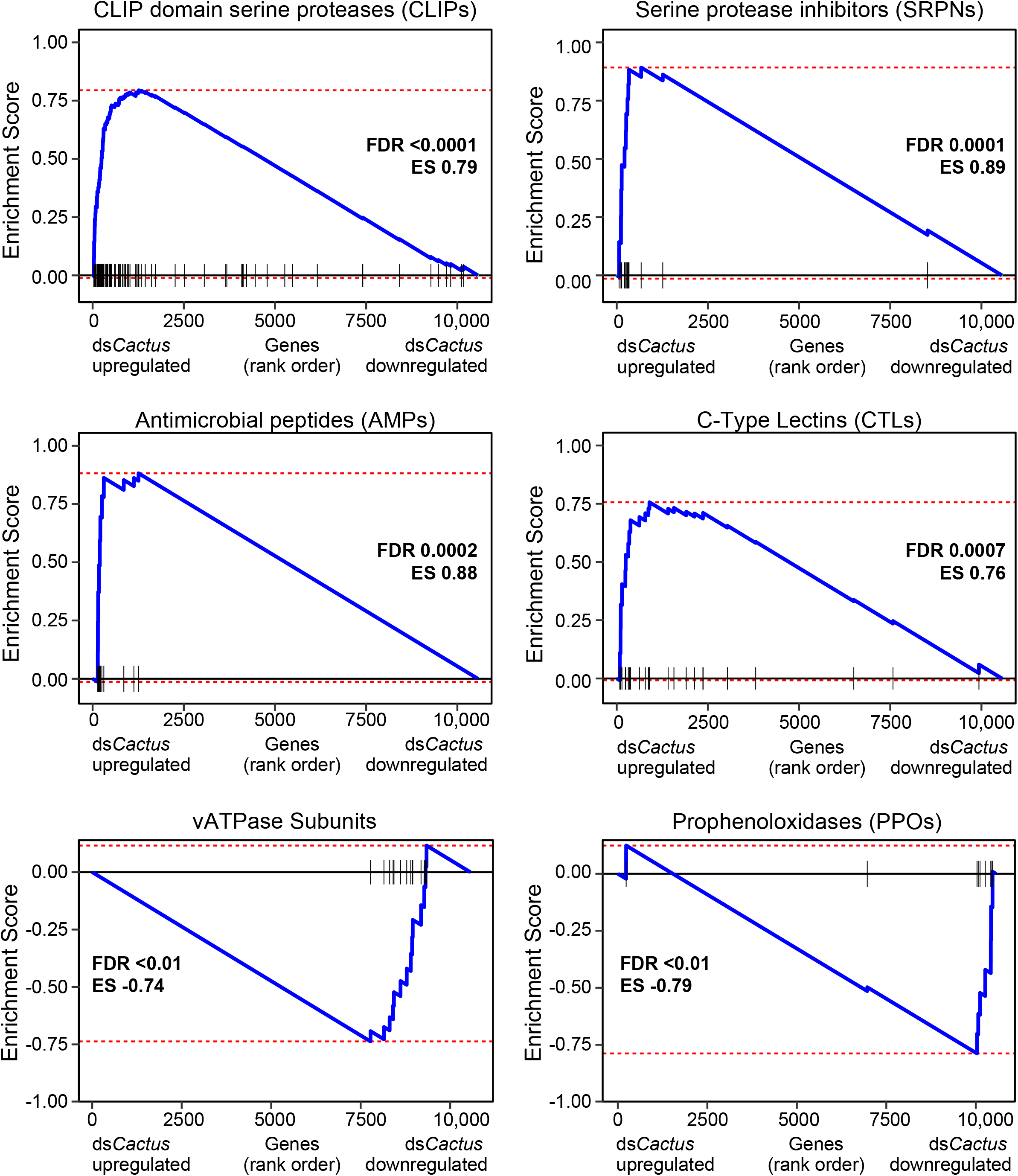
Gene Set Enrichment Analysis of Malpighian tubule transcriptional responses to Toll activation. GSEA was performed with six manually curated gene sets which showed significant associations in our Malpighian tubule transcriptomic comparisons. Each panel represents the association of one gene set (CLIPs, SRPNs, AMPs, CTLs, vATPases, and PPOs) where the y-axis shows the Enrichment Score (ES), and the x-axis shows the log_2_FC rank of all genes detected in the transcriptome. Black ticks represent where genes in the respective gene list fall on the continuum of ranked genes. ES and the corrected P-value (FDR) for each gene set shown. ES scores indicate genes enriched in those that are upregulated (positive ES) or downregulated (negative ES) following ds*Cactus* treatment. Full statistical description of all lists can be found in Table S5.

### Comparing the hemolymph proteome of control mosquitoes and dsCactus-treated mosquitoes using high-resolution LC-MS/MS

Mass spectrometry-based proteomic analysis of hemolymph extracted from control and ds*Cactus*-treated mosquitoes revealed a total of 1,319 proteins corresponding to 14,018 peptides. The complete list of peptides and their corresponding protein identifications is provided in the supplemental information (Table S5, Table S6). This is the first comprehensive hemolymph proteomic profiling of adult *Ae. aegypti* mosquitoes using nano-LC coupled to high-resolution mass spectrometry. We performed label-free quantification (LFQ) analysis to identify differentially expressed hemolymph proteins between control and ds*Cactus*-treated mosquitoes. Among the hemolymph samples, approximately 64% of the variability in the data is likely related to treatment with ds*Cactus* as represented by PCA (Figure 4A). The Pearson correlation coefficients among the replicates fall within the range of 0.85-0.99, showing a close association between the replicates and suggesting high repeatability (Figure S2).

**Figure 4:**
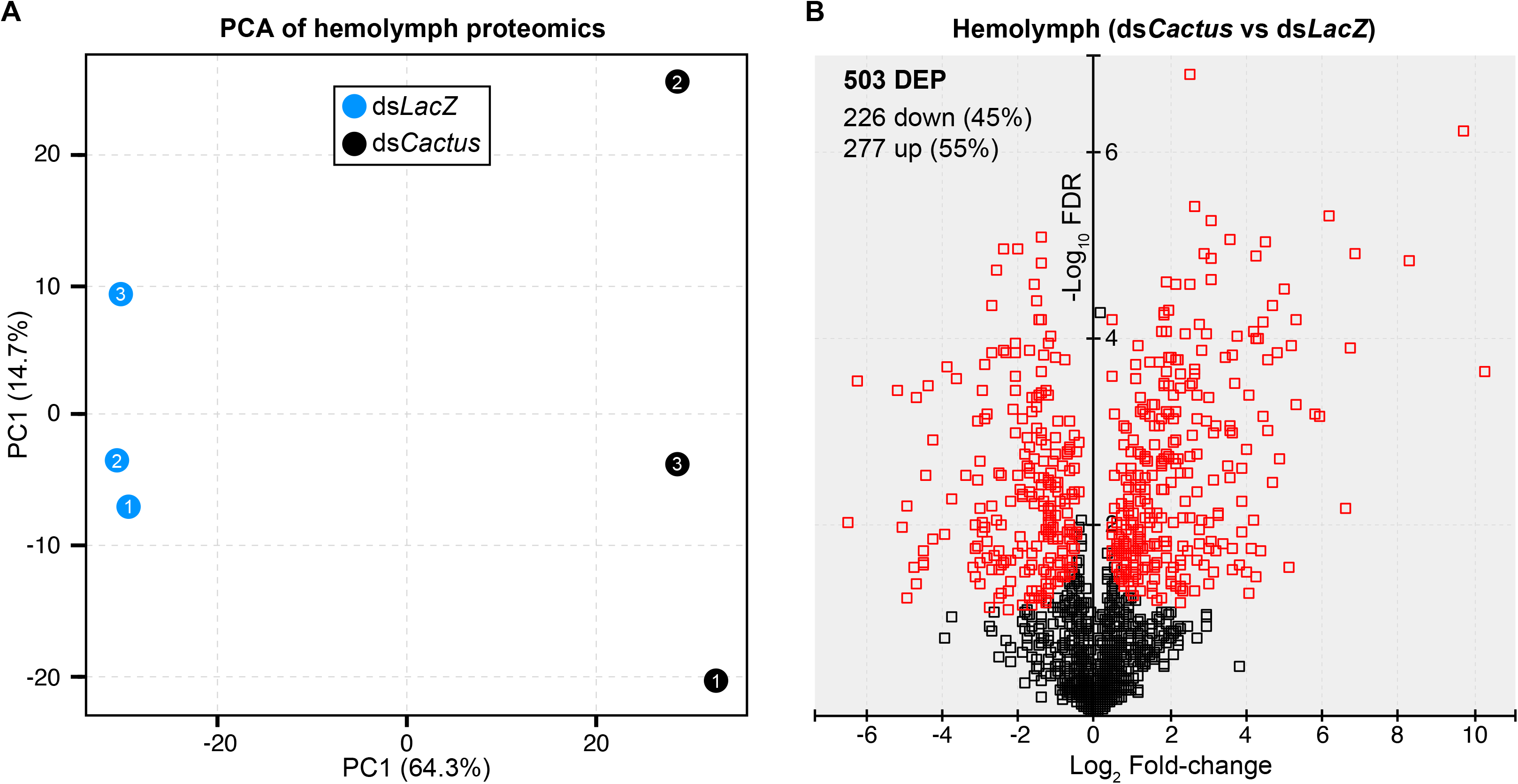
Proteomic analysis of hemolymph following Toll activation. (**A**) Principal component analysis of all six hemolymph samples to show global variation; clustering along the first principal component was observed between ds*LacZ* (blue circles) and ds*Cactus* (black circles) treatments. (**B**) A volcano plot was generated to estimate protein level changes among proteins identified. Red symbols indicate significantly differentially expressed proteins.

To determine differentially expressed proteins between the hemolymph proteomes of control and *dsCactus-*treated mosquitoes, the normalized LFQ intensities of the identified peptides between the two experimental groups were compared. The filtered analysis led to the identification of 277 significantly upregulated and 226 significantly downregulated proteins (Table S7). Protein set enrichment analysis (PSEA) using fold change values from this analysis showed an enrichment of CLIPs, SRPNs, CTLs and other proteases, indicating the possibility of downstream melanization effector mechanisms in the hemolymph of *dsCactus*-treated mosquitoes (Figure S3, Table S4). Interestingly, the enrichment of CLIPs, which includes one of the most highly enriched proteins AAEL022822, along with the increased abundance of TEP20 in the hemolymph suggests the possible accumulation of putative complement pathway components following *dsCactus*-treatment. This pathway has not yet been characterized in *Ae. aegypti*, but it plays a prominent role in anti-plasmodial, anti-bacterial, and anti-fungal immunity in *An. gambiae* [24, 33, 36, 37, 38, 39, 40]. Taken together, our data suggests that *dsCactus*-treatment increases expression of proteins involved in mosquito immunity with specific enrichment of proteins involved in the Toll and putative complement pathways.

### Determining the potential for a Malpighian-tubule specific contribution to systemic humoral immunity

Lastly, we tested the hypothesis of whether the hemolymph proteins upregulated in *dsCactus*-treatment might have an origin in the MT. Using GSEA, we observed that proteins significantly increased in the *dsCactus* hemolymph tended to be similarly enriched in the *dsCactus* MT RNA-seq dataset (Figure 5A). We then identified 168 transcripts which were significantly upregulated by *dsCactus*-treatment specifically in the MT but not in the whole body (Figure 2B). When this list was intersected with hemolymph proteins upregulated by *dsCactus*-treatment (Figure 5B) we found a statistically significant overlap of 39 transcripts/proteins (Representation factor 10.0; p < 1.868e-28 for tubules). This group contained several putatively secreted Toll and complement related genes such as several CLIPs, SRPN3, and TEP20 (Table S8). While this does not definitively confirm that these proteins are secreted from the Malpighian tubules into the hemolymph following *dsCactus*-treatment, we suggest this as a hypothesis.

**Figure 5:**
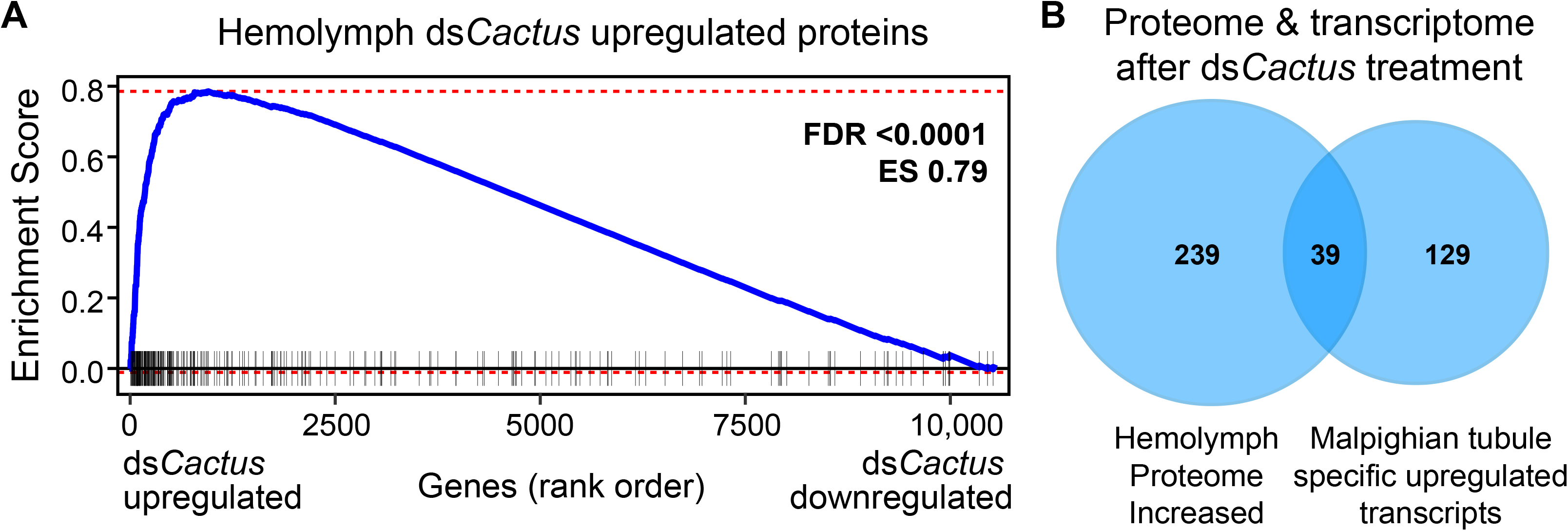
Comparison of Malpighian tubule transcriptomic and hemolymph proteomic responses following Toll activation. (**A**) GSEA was performed on the Malpighian tubule RNA-seq using a gene set consisting of proteins that were significantly upregulated in the hemolymph proteomics. The y-axis shows the Enrichment Score (ES), and the x-axis shows the log_2_FC rank of all genes detected in the Malpighian tubule transcriptome. Black ticks represent where proteins in the hemolymph upregulated proteome gene list fall on the continuum of ranked genes. ES and the corrected P-value (FDR) for this analysis is shown. (**B**) Among the 168 genes that were significantly upregulated only in the Malpighian tubules upon ds*Cactus* treatment, 39 were also upregulated in the hemolymph proteome.

## Discussion

Our study demonstrates the direct transcriptional response of MT to *Cactus* systemic silencing and their capacity to broadly upregulate Toll pathway target genes in *Ae. aegypti*. Extensive work in *Drosophila* and different mosquito species has shown that hyperactivation of Toll-signaling using a variety of approaches results in increased immune activation and refractoriness to immune challenge [7, 23, 26, 27, 33, 34, 41, 42, 43]. Studies in *Ae. aegypti* have shown that *dsCactus*-treatment drives constitutive Toll-signaling, immune activation, and increased refractoriness to viral [44], fungal [41], and filarial nematode infections [7]. In *An. gambiae*, activation of Toll-signaling mediated by *dsCactus*-treatment resulted in increased basal expression of TEP1 and LRIM1, immune genes that inhibit development of *Plasmodium*, resulting in lower burden of parasite infection [33]. Similar studies using transgenic strains of *Ae. aegypti* overexpressing Rel1, the NF-κB transcription factor activated downstream of Toll-signaling, showed increased expression of immune genes in the absence of a pathogenic challenge [41]. The significant degree of correlation between other transcriptomes profiled during Toll activation (e.g. REL1 overexpression; infection etc.) and our dataset (Table S4) confirm earlier findings that silencing of the *Cactus* gene activates the Toll pathway.

This study extends our current understanding of Toll pathway signaling, by demonstrating the ability of the MT to undergo a Toll-based immune response. Our previous transcriptomic studies comparing mosquitoes susceptible and refractory to the filarial nematode *D. immitis* suggested that Toll pathway activation was sufficient to block the worm development within the tubule [7]. Correlation of the genes upregulated in *dsCactus*-treatment samples between the MT and whole body (Figure 2B) along with the specific upregulation of canonical Toll targets such as CTLs, CLIPs, SRPNs, and PPOs in the MT (Figure 3) strongly suggests that the Malpighian tubules themselves generate immune factors for pathogen defense. Interestingly, subunits of the vATPase were also downregulated following *dsCactus*-treatment (Figure 3). vATPase is a proton (H+) pump that has been shown in *Ae. aegypti* to be responsible for the bulk of Malpighian tubule transepithelial secretion of KCl and NaCl. The location of vATPase in Malpighian tubules is in the apical brush border membrane of principal cells and it is responsible for mediating a substantial part of post-blood meal diuresis. It was previously shown that vATPases were globally downregulated in Ae. albopictus MT following blood feeding [45]. To date, there has been no evidence in mosquitoes of a direct link between vATPase expression and immunity. However, work in *Drosophila* suggests there is an important connection between physiological desiccation stress (metabolism) and immunity [46, 47, 48, 49], so it stands a similar pathway may be at work in mosquitoes although further studies are needed to confirm this relationship.

Even more intriguing is the possibility that the tubules play a more systemic role in Toll-based immunity. The relative contribution of the MT to the immune protein milieu in the hemolymph of mosquitoes has not been defined and we only sought to begin to explore this hypothesis in this study. We found a subset of genes that were enriched during Toll activation specifically in the MT that encode proteins that were also significantly increased in the hemolymph under the same conditions (Table S8). This suggests the possibility that such proteins are derived from the tubules. There is precedent for a significant systemic immune contribution from the MT in *Drosophila* as antibacterial immunity develops in larvae only after development of the MT [50]. The MT are also strictly controlled by neuroendocrine factors, and in adult flies suffering from desiccation stress, ecdysone is locally produced in MT to increase expression of immune genes and increase host defense [46]. Although the numbers are small, sessile hemocytes are present attached to the MT [51]. Additionally, blocking the Malpighian tubule immune response immunocompromises flies even when the fat body is intact, indicating that the contribution of the MT is systemically important [52, 53]. Future work and new genetic tools will be needed to establish the functional contribution of MT to systemic immunity in mosquitoes.

## Conclusions

This work has characterized the transcriptome of MT from mosquitoes treated with *dsCactus* RNA and shown for the first time that the MT of *Ae. aegypti* can activate Toll-pathway immune genes locally in response. We have also profiled the hemolymph proteins of Toll-activated mosquitoes and identified a subset of proteins present whose transcripts are differentially upregulated in the MT, which suggests that possibility that the MT can contribute substantially to humoral immunity. Collectively, this work contributes to the growing body of evidence that the MT are an essential immune tissue with similarly active pathways to other known immune tissues (e.g. fatbody and hemocytes), but possibly unique physiological roles, in the mosquito immune response.

## Abbreviations

MT: Malpighian tubules
RNAi: RNA interference
KD: knockdown
SRPN: Serine Protease Inhibitor
CTL: C-Type Lectin
CLIP: CLIP-domain serine protease
LRIM: Leucine-rich repeat Immune Protein
TEP: Thioester containing protein
PPO: Prophenoloxidase
AMP: Anti-microbial peptide
vATPase: vacuolar-type ATPase
GSEA: Gene Set Enrichment Analysis
PSEA: Protein Set Enrichment Analysis
PCA: Principal Component Analysis
MS: Mass-spectrometry
LFQ: Label-free quantitative MS

## Declarations

## Acknowledgements

We thank Abigail R. McCrea and Sarah Dysinger for technical assistance producing mosquitoes and other materials for this study.

## Availability of data and materials

All scripts used to analyze transcriptomic data were written in R or bash and are publicly available on Rpubs (https://rpubs.com/shanedenecke/Cactus_KD_analysis). Raw mRNA sequencing reads are available on NCBI Sequence Read Archive (PRJNA758024). Our data meet all the standards regarding the Minimum Information About a Proteomics Experiment (MIAPE), and data have been deposited to the ProteomeXchange Consortium (http://www.proteomexchange.org) via the PRIDE partner repository (px-submission #607630).

## Authors’ contributions

SDS, SBD, SD, and MP Conceived and designed the analysis; CD and MP Collected the data; SDS, SBD, SD Performed the analysis; SDS, SBD, SD, and MP Wrote the paper.

## Ethics approval and consent to participate

Not applicable

## Consent for publication

Not applicable

## Figure Legends

**Figure S1: Gene Set Enrichment Analysis of Malpighian tubule transcriptional responses to Toll activation**. GSEA was performed with four manually curated gene sets from previously published transcriptomic studies which showed significant associations in our Malpighian tubule transcriptomic comparisons. Each panel represents the association of one gene set where the y-axis shows the Enrichment Score (ES), and the x-axis shows the log_2_FC rank of all genes detected in the transcriptome. Black ticks represent where genes in the respective gene list fall on the continuum of ranked genes. ES and the corrected P-value (FDR) for each gene set shown. ES scores indicate genes are enriched in those that are upregulated (positive ES) or downregulated (negative ES) following ds*Cactus* treatment. Full statistical description of all lists can be found in Table S5.

**Figure S2: Hemolymph proteomic replicates are highly correlated**. Correlation analysis to demonstrate consistency between hemolymph mass spectrometry proteomic replicates. Replicate 1 (x-axis) is shown against replicate 3 (y-axis) with the correlation coefficient ρ value (0.94) shown on the graph. Below the graph are the ρ values for the other replicate comparisons, which are also highly correlated to each other.

**Figure S3: Protein Set Enrichment Analysis of hemolymph response to Toll activation**. The y-axis shows the Enrichment Score (ES) and the x-axis shows the log_2_FC rank of all proteins detected in the hemolymph proteome. Black ticks represent where genes of the respective proteins from curated lists fall on the continuum of ranked proteins. Full statistical description of all lists can be found in Table S5.

## Table legends

**Table S1**. Gene lists used for the GSEA and PSEA. Column headings are gene list names and rows beneath are the VectorBase gene identifiers belong to each list. Some lists are manually curated and others are derived from previously published data sets.

**Table S2**. Overview of Malpighian tubules and whole mosquito mRNA sequencing. An overview of the transcriptomic results is shown including total reads, mapping rates, and the percent of genes mapping to known protein encoding genes.

**Table S3**. Differentially expressed genes from Malpighian tubules and whole mosquito mRNA sequencing. All RNA-seq runs are shown along with their corresponding metadata. Down- and up-regulated genes in MT are shown in grey and blue shading, respectively. Down- and up-regulated genes in whole mosquitoes are shown in orange and green shading, respectively.

**Table S4**. Statistical values for the transcriptomic GSEA and PSEA. Gene and protein sets enriched in MT transcriptomics and hemolymph proteomics.

**Table S5**. Peptide-level output of the hemolymph proteomics.

**Table S6**. Proteins detected in hemolymph proteomics with their associated metadata.

**Table S7**. Differentially expressed proteins in hemolymph proteomics from ds*Cactus* treated mosquitoes.

**Table S8**. Intersection of the genes which were commonly found between i) genes upregulated in the ds*Cactus* treated MT but not upregulated in the ds*Cactus* treated whole body samples and ii) proteins upregulated in the ds*Cactus* treated hemolymph proteome.

