## Supplementary figures and images for "*Aedes aegypti* Malpighian tubules are immunologically activated following systemic Toll activation"

### Figure S1

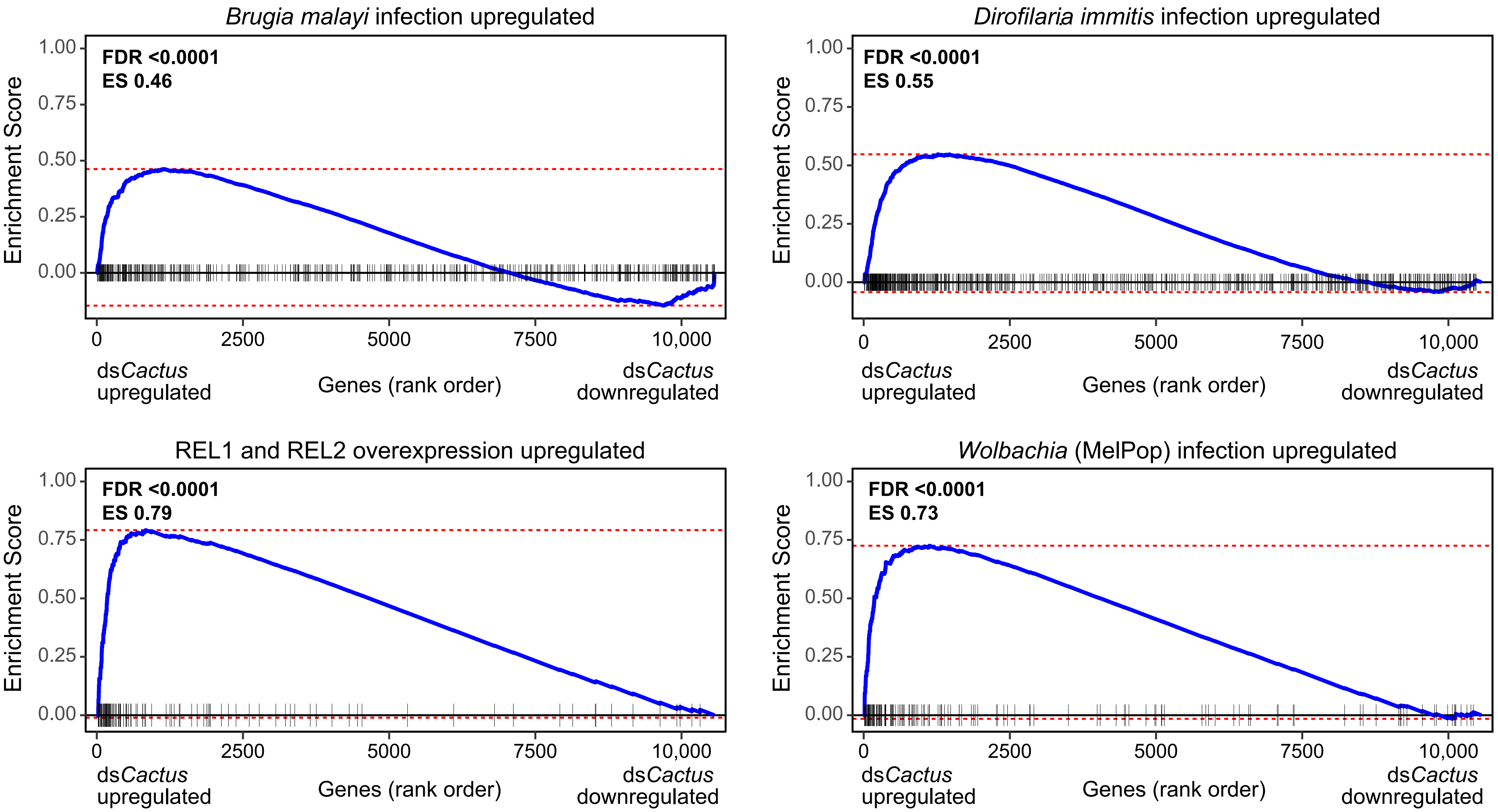

### Figure S2

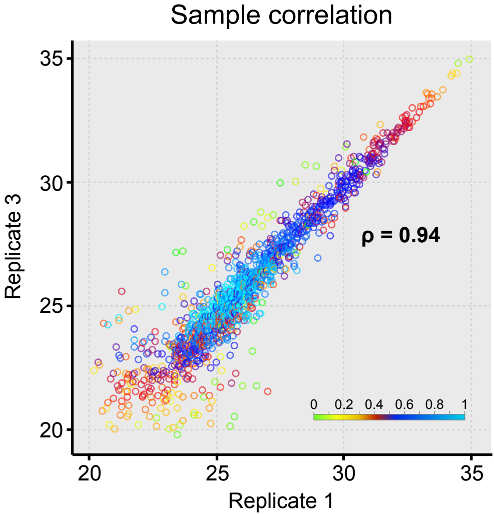


Replicate 1 vs 2 (ρ = 0.91)
Replicate 2 vs 3 (ρ = 0.93)

**Figure S2**

### Figure S3

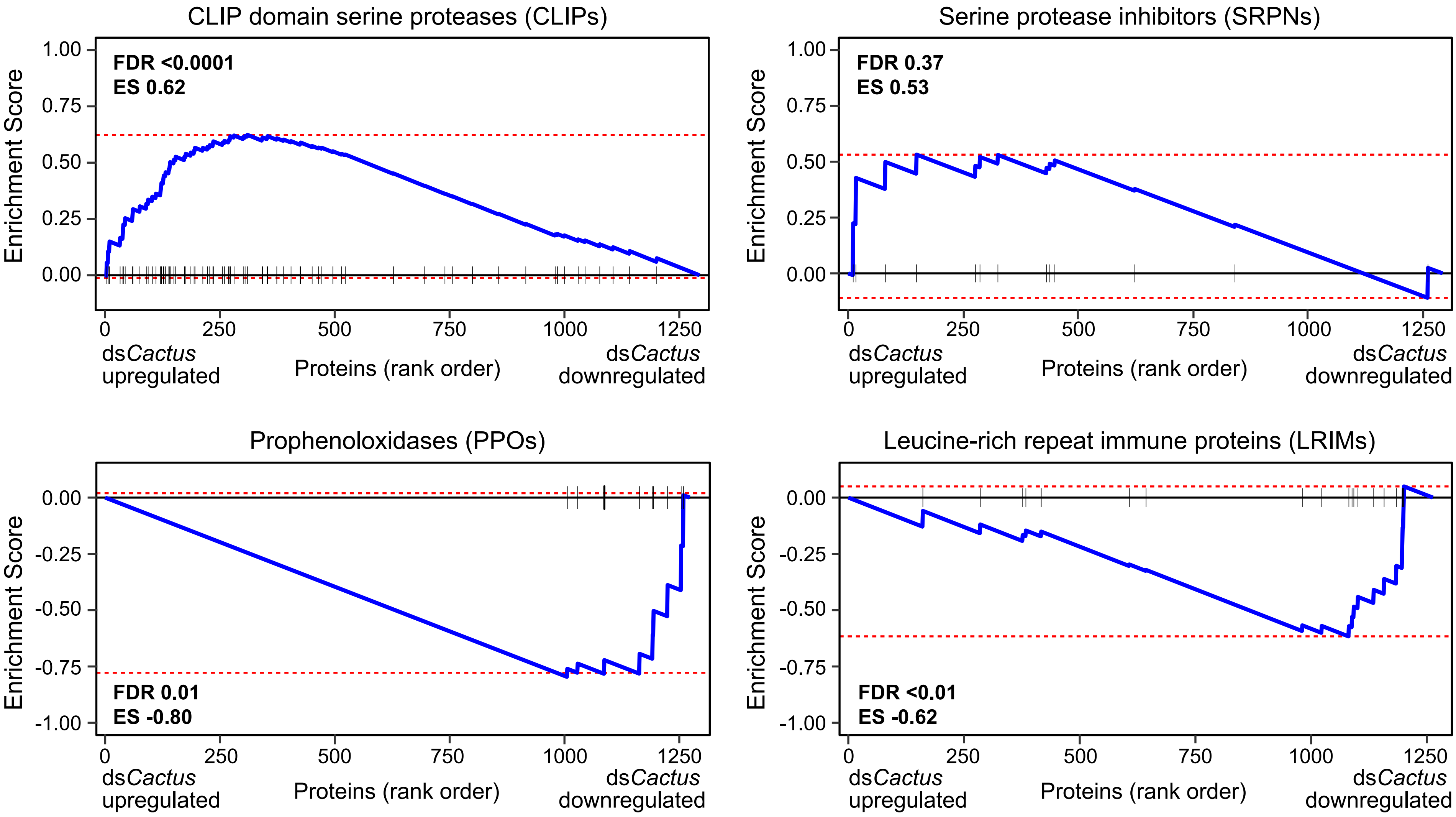
